# Engineering developmental transitions during human neurulation through modular integration of multimodal spatial cues

**DOI:** 10.64898/2026.10.07.757222

**Authors:** Masaya Hagiwara, Sae Nozaki, Yosuke Sugino, Tomohiro Mimura, Yen Xuan Ngo, Laura Yuriko Gonzalez-Teshima, Isabel Koh, Yasuhiro Inoue, Isao Matsuo

**Affiliations:** Centre for Biosystems Dynamics Research, RIKEN, Kobe, 650-0047, Japan; Department of Micro Engineering, Graduate School of Engineering, Kyoto University, Kyoto, 615-8540, Japan; Centre for Brain Science, RIKEN, Wako, 351-0106, Japan; Department of Molecular Embryology, Research Institute, Osaka Women’s and Children’s Hospital, Osaka Prefectural Hospital Organization, Izumi Osaka 594-1101, Japan

## Abstract

Reconstituting developmental transitions *in vitro* requires reproducing complex spatial cues that guide cellular self-organization. Here, we developed MoSAIC (Modular Spatial Assembly and Integration of Cues), a modular platform that reconstructs multimodal mechano-chemical spatial cues by decomposing complex culture operations and subsequently integrating them. Central to MoSAIC is a composite architecture combining rigid polycarbonate structures with soft extracellular matrix hydrogels, providing geometric precision and structural stability while permitting cellular self-organization and tissue deformation. Using human induced pluripotent stem cells, MoSAIC enabled independent control of tissue geometry, ECM localization, physical boundaries, and morphogen gradients, together with defined mechanical and chemical perturbations. Their integration guided a continuous ectodermal sheet through neural induction, bending, and bifurcation into neural and epidermal tissues, followed by polarized neural tube formation and dorsoventral patterning. The resulting tissue exhibited posterior identity, while non-genetic perturbations enabled interrogation of mechanisms governing neural tube closure.

## Main

Development is not merely a process of cell fate specification, but a dynamic sequence of tissue transformations through which cells interpret positional information from their surrounding environment and collectively construct functional architectures^1^. Spatial information encoded in tissue geometry, extracellular matrix organization and soluble morphogen distribution coordinates cell differentiation, tissue deformation and multicellular self-organization^2^. Such environmental cues are particularly critical during neurulation, in which an initially continuous ectodermal sheet is largely specified into neural and non-neural territories, undergoes bending and closure, and ultimately separates into two distinct tissues: the neural tube and the surface ectoderm^3^. Thus, neural tube formation represents a paradigmatic developmental transition in which spatially organized biochemical and mechanical signals drive the bifurcation of a single epithelial tissue into distinct anatomical structures.

Stem cell-derived organoids have emerged as powerful models for studying human development and disease^4,5^. However, most organoid systems are generated by culturing dissociated or aggregated cells under spatially uniform conditions and therefore rely predominantly on intrinsic self-organization^6^. Although such systems can reproduce aspects of cell fate specification and regional patterning, they often form spherical structures with heterogeneous domains rather than undergoing the directed tissue-level transformations observed in embryos^5, 7^. Recent advances in patterned culture substrates^8^, microfluidic morphogen delivery^9,10^, Assembloid^11-13^, bioprinter^14,15^ and synthetic hydrogel technologies^16-18^ have enabled the induction of regionalized cell identities and interactions between preformed tissue modules. Nevertheless, these approaches have largely focused on reproducing endpoint patterning rather than the dynamic transitions between developmental states^5, 19^. In particular, an *in vitro* system that reconstructs tissue bifurcation, in which a single continuous tissue autonomously separates into two distinct tissue identities through morphogenetic transformation, has barely achieved.

Reconstructing such developmental transitions is important not only for understanding human morphogenesis, but also for modelling congenital disorders that arise from failures in developmental processes themselves. Neural tube defects, for example, are caused by disruptions in the coordinated sequence of patterning, epithelial remodelling, tissue bending and closure, rather than by abnormalities in terminal cell fate alone^3,20,21^. Therefore, models that only reproduce final tissue identities are insufficient to capture the developmental mechanisms underlying these human disorders. To investigate how tissue form emerges, fails or can be controlled, it is necessary to rebuild the spatiotemporal environmental cues that guide morphogenesis, including both biochemical and mechanical inputs.

A major obstacle to this goal is the lack of experimental platforms capable of integrating multimodal spatial information in a controllable and practical manner. Developmental morphogenesis is orchestrated by the combined effects of mechanical and chemical interactions associated from the environmental factors such as tissue geometry, extracellular matrix organization and soluble signalling gradients^1^ (Fig. 1a). However, simultaneous control of these parameters remains technically challenging. Assembloid approaches primarily manipulate the spatial arrangement of cell populations, whereas microfluidic systems are well suited for soluble factor delivery but rarely provide integrated control over tissue geometry, extracellular matrix architecture and spatio-temporal soluble factors. Increasing the complexity of culture systems can also reduce reproducibility, accessibility and experimental throughput. Thus, the inability to simultaneously reconstruct the multimodal cellular microenvironment remains a central bottleneck in engineering organoids that mimic developmental processes rather than static developmental outcomes.

**Figure 1.**
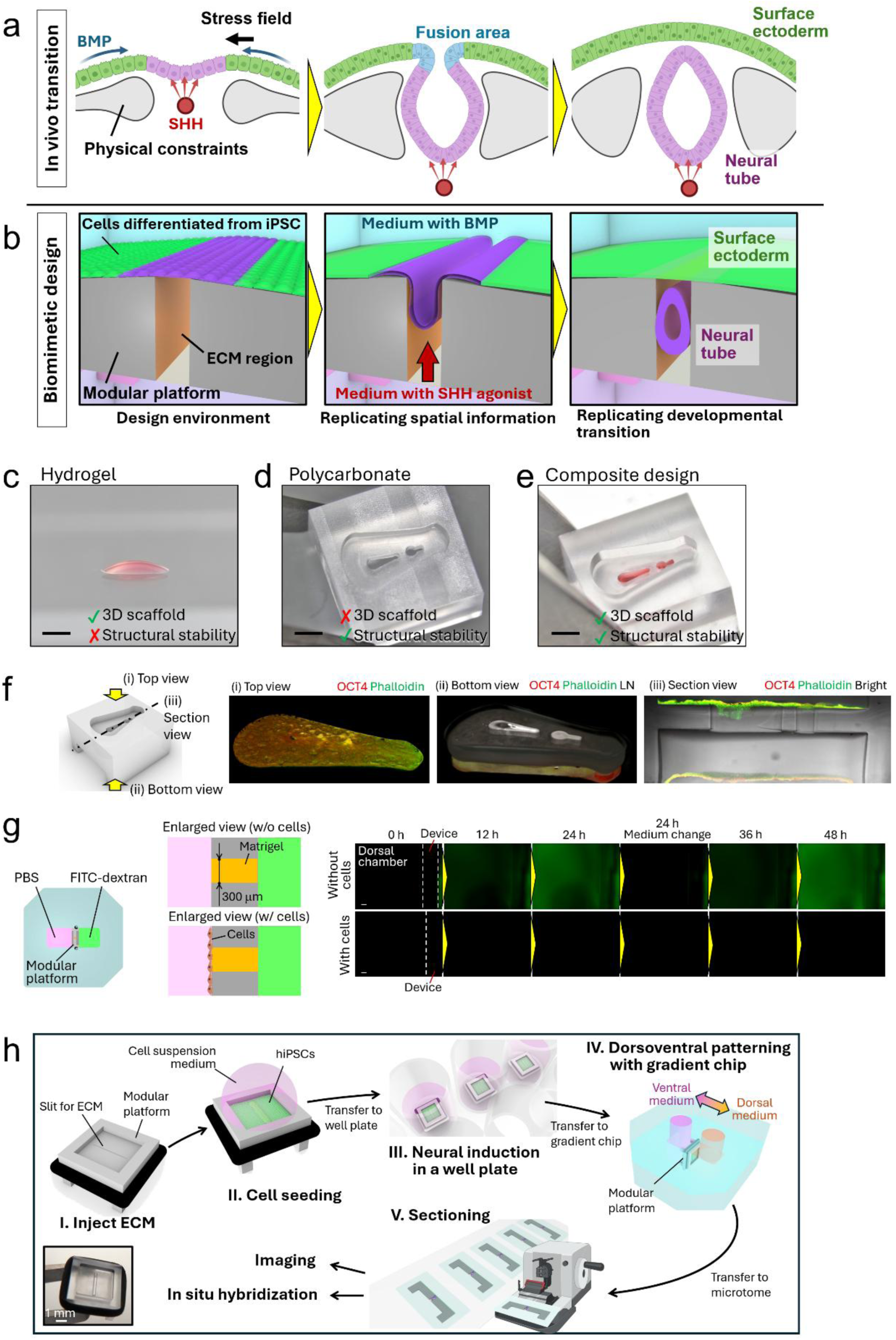
Development of the MoSAIC platform for reconstructing multimodal spatial cues during organ development. **a,** Schematic illustration of the spatially coordinated mechanical and biochemical cues involved in neural tube development *in vivo*. **b,** Conceptual illustration highlighting the importance of engineering biomimetic microenvironments with multimodal spatial cues to recapitulate organ development *in vitro*. **c,** Soft extracellular matrix (ECM) hydrogels provide a permissive three-dimensional scaffold for cell growth and tissue morphogenesis but exhibit limited structural stability. **d,** Rigid polycarbonate structures can be fabricated with precisely defined three-dimensional geometries and provide mechanical stability but lack the biological functionality required to support three-dimensional tissue development. **e,** Composite design integrating soft ECM hydrogels with rigid polycarbonate structures combines a permissive biological microenvironment with geometric precision and structural stability. The polycarbonate framework enables ECM hydrogels to be spatially confined within predefined geometries. Scale bar: 1 mm. **f,** Demonstration of spatial control over ECM localization and cell seeding geometry, together with the flexibility for multidirectional imaging. Matrigel was localized within predefined regions of the polycarbonate device, and hiPSCs were seeded on both sides of the device. **g,** Demonstration of spatially controlled molecular transport following integration of the modular device into a microfluidic chip. FITC-labelled dextran solution was introduced into the right chamber, while PBS was introduced into the left chamber. Solutions were replaced every 24 h to maintain a molecular concentration gradient across the ECM within the modular device. Scale bar: 200 μm. **h,** Schematic illustration of the MoSAIC concept, in which individual processes for spatial cue control are modularized and subsequently integrated to reconstruct multimodal microenvironments. The platform also preserves the spatial organization of cells and ECM during downstream analyses.

Here, we developed MoSAIC (Modular Spatial Assembly and Integration of Cues), a modular platform that enables the integration of multimodal spatial cues to reconstruct developmental transitions during neurulation. The soft materials such as cells and ECM can be modularized and handled with a pair of tweezers by the device made by polycarbonate. This modularity allows us to divide the control process into the simple operations and sum of the simple operations can recapitulate complex spatial cues generated *in vivo* (Fig. 1b). By independently controlling cell seeding geometry and size, extracellular matrix organization and soluble morphogen distribution, we establish a human iPSC-derived neural tube morphogenesis model that recapitulates the transition from a single undifferentiated ectodermal sheet to separated neural and epidermal tissues. Importantly, this system reproduces the spontaneous segregation of an initially continuous epithelial sheet into distinct tissue domains, thereby mimicking tissue bifurcation during neurulation. Modularity of the MoSAIC also enables us to pass the developed neural tube organoid to the post-experimental analysis retaining spatial information after sectioning the developed organoid together with the polycarbonate frame for immunostaining and in situ hybridization.

Our findings provide a framework for engineering developmental morphogenesis as a programmable process and demonstrate that organoid systems can move beyond static fate patterning toward controllable tissue dynamics.

## Results

### Development of MoSAIC system

A major challenge in engineering three-dimensional culture environments is the mechanical fragility and structural instability of extracellular matrix (ECM) hydrogels, despite their essential role in supporting cell growth and tissue morphogenesis (Fig. 1c). In contrast, rigid materials such as polycarbonate provide structural stability and precisely defined geometries (Fig. 1d). We therefore combined soft ECM hydrogels with a rigid polycarbonate frame to integrate complementary functions: the hydrogel provides a permissive biological microenvironment, whereas the rigid structure defines and preserves its spatial organization. Using this composite design, ECM hydrogels could be localized within predefined regions of the polycarbonate device through surface tension and retained after gelation (Fig. 1e). The surrounding structure also defined the cell-seeding geometry, enabling independent spatial organization of cells and ECM. Importantly, the rigid frame could be manipulated with tweezers, allowing the construct to be flipped for imaging and subsequently sectioned together with the biological sample. Figure 1f demonstrates these capabilities: Matrigel was localized within the openings and along the backside of the device, while hiPSCs were seeded onto both surfaces. The construct could be imaged from either orientation, and cross-sectional images were obtained by sectioning the cells, ECM, and polycarbonate together. Thus, the composite architecture enabled fragile cell–ECM constructs to be spatially organized, manipulated, and analysed while preserving their original configuration.

The modular architecture further enabled integration with microfluidic systems for spatial control of soluble factors. To examine molecular transport, a device containing a 300 µm-wide Matrigel-filled slit was incorporated into a fluidic chip (Fig. 1g). When 20-kDa FITC-labelled dextran was introduced into one chamber, fluorescence progressively appeared in the opposite chamber through diffusion across the hydrogel. The fluorescence signal increased over time and was reset following medium exchange at 24 h. This diffusion-mediated transport enables concentration gradients to be established across the ECM, consistent with our previously developed gradient systems^22^. In contrast, when a continuous cell layer covered the Matrigel region, dextran transport into the opposite chamber was restricted, indicating that soluble factors could be delivered locally to defined regions of the cell layer. These results demonstrate the integration of ECM localization and spatially controlled molecular delivery within the same device.

Together, these features establish the material and operational basis of MoSAIC (Modular Spatial Assembly and Integration of Cues). By combining the structural precision of rigid polycarbonate with the biological permissiveness of soft ECM hydrogels, MoSAIC enables cells, ECM, and soluble factors to be independently organized and subsequently integrated as multimodal spatial cues. Its modular architecture decomposes complex culture operations into manageable steps, including ECM patterning, cell positioning, molecular delivery, transfer between culture environments, imaging, and sectioning, before integrating them into a defined experimental workflow.

We next applied MoSAIC to reconstruct developmental transitions during neural tube formation (Fig. 1h). A slit within the polycarbonate frame confines ECM through surface tension, while surrounding walls define the geometry and dimensions of the cell-seeding region. Following cell seeding, the construct is initially cultured in a conventional well plate for neural induction and subsequently transferred into a fluidic chip, where distinct media are supplied from opposite sides to establish dorsal and ventral signalling environments (Supplementary Video 1). Localized diffusion through the ECM-filled slit further enables soluble-factor gradients, mimicking spatially restricted developmental cues such as notochord-derived SHH. After culture, the entire construct can be sectioned for spatially resolved analyses, including immunostaining and in situ hybridization. Thus, MoSAIC integrates the construction, manipulation, perturbation, and analysis of spatially organized tissues within a single modular framework for engineering developmental microenvironments.

### Designing MoSAIC

MoSAIC can therefore be configured to reconstruct multiple spatial features associated with neurulation. However, the large number of possible device configurations necessitated the identification of design parameters that support neural tube morphogenesis. Neurulation involves spatially coordinated interactions among the neural plate, surrounding surface ectoderm, and ECM. Because ECM provides both biochemical and mechanical cues and is spatially heterogeneous around the neural plate *in vivo*, we first asked whether localized ECM could be used to define the spatial organization of neural and non-neural ectodermal domains.

To examine the effect of spatial ECM control during neurulation-associated patterning, Matrigel was patterned with widths of 250 µm or 500 µm on a glass bottom dish respectively, and iPSCs were seeded both on and around the Matrigel region. After 3 days of neural induction followed by 2 days of culture in dorsal medium containing recombinant BMP4 and the WNT inhibitor IWP2, the cells developed spatially segregated expression patterns. N-cadherin-positive cells preferentially emerged within the Matrigel-patterned region, whereas E-cadherin-positive cells occupied the surrounding area (Fig. 2a and Supplementary Fig. 1a). Quantification of E-cadherin and N-cadherin distribution confirmed that ECM localization was sufficient to spatially organize two distinct ectodermal domains corresponding to the neural plate and surrounding surface ectoderm (Fig. 2b and Supplementary Fig. 1b). These results established ECM localization as a key design parameter for defining the tissue domains required for subsequent neural tube morphogenesis.

**Figure 2.**
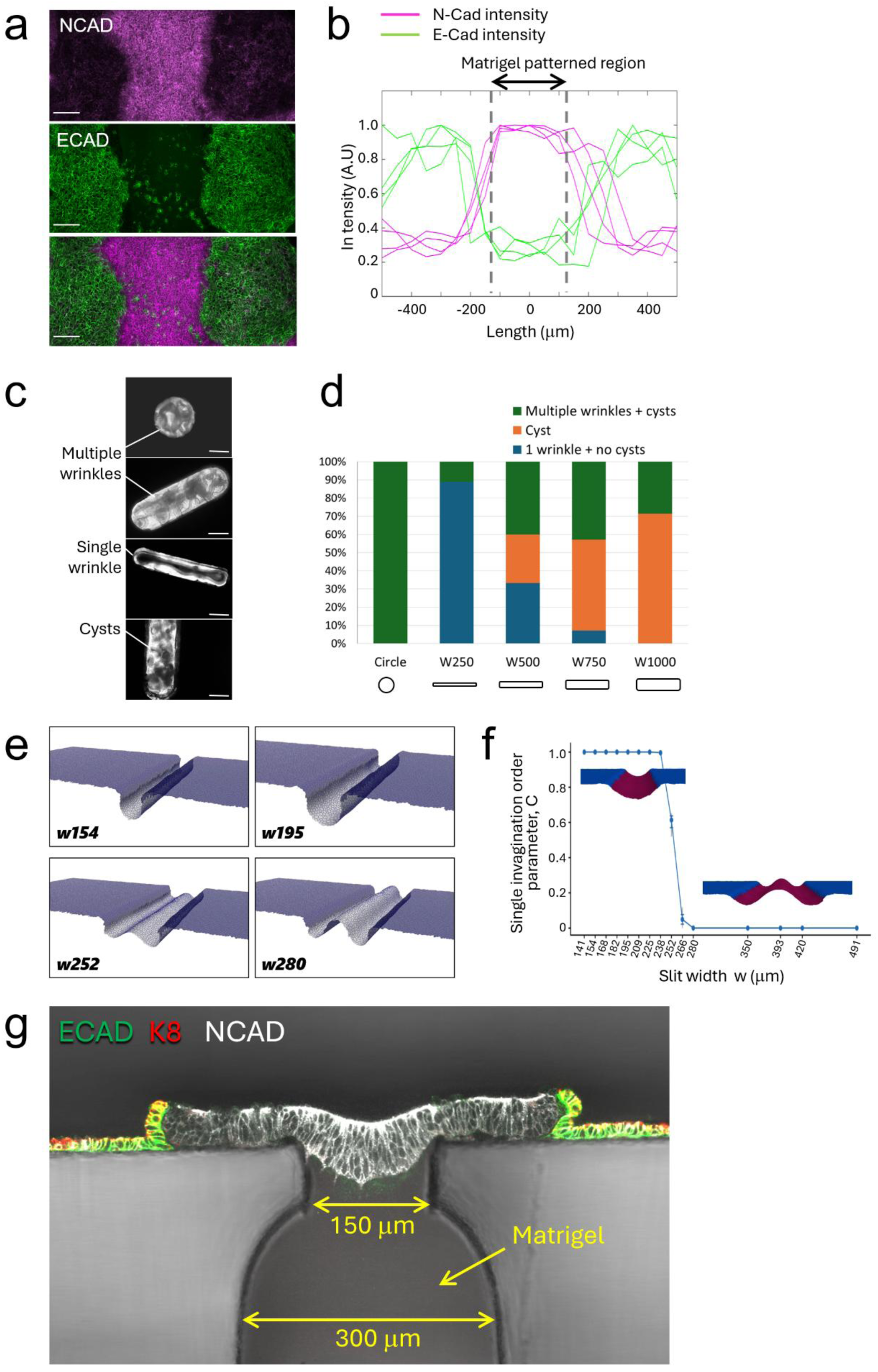
Design of the MoSAIC system to recapitulate developmental transitions during neurulation. **a,** Effect of spatially localized Matrigel on hiPSC differentiation. Matrigel was patterned into a rectangular region with a width of 250 μm. N-cadherin-positive cells emerged within the Matrigel-patterned region, whereas cells outside this region predominantly expressed E-cadherin. Scale bar, 100 μm. **b,** Quantification of the spatial distribution of E-cadherin- and N-cadherin-positive cells in the presence of a 250-μm-wide rectangular Matrigel pattern (n = 4). The dot lines indicate the boundary of the Matrigel region. **c,** Representative tissue morphologies generated using Matrigel-filled slits with different widths and geometries. Scale bar, 500 μm. **d,** Quantification of morphological outcomes as a function of slit geometry and width. **e,** Mathematical simulations illustrating the effects of slit width on tissue morphogenesis. **f,** Transition from single-wave to double-wave deformation patterns with increasing slit width. The single-invagination order parameter, C, characterizes the transition in the number of deformation waves. **g,** Final configuration of the MoSAIC platform designed to recapitulate developmental transitions during neurulation.

Although ECM patterning enabled spatial segregation of neural and epidermal identities, the geometry and dimensions of the neural region were expected to critically influence its subsequent morphogenesis including closure process. When iPSCs were seeded on a dish and apply a neural induction medium, many of wrinkle patterns emerged (Supplementary Fig. 1c and Video 2). This morphology needs to be guided properly to make consistent tube formation for neural tube. We therefore fabricated a series of polycarbonate devices with different slit geometries and dimensions to systematically examine how the shape and size of the Matrigel-defined neural region affected morphogenesis. During culture in neural induction medium, hiPSC-derived cell sheets underwent spontaneous deformation, generating wrinkles or cystic structures depending on the geometry and dimensions of the patterned region (Fig. 2c). A narrow rectangular pattern 250 µm in width most frequently resulted in the formation of a single pair of longitudinal folds, which subsequently developed into a tubular structure. In contrast, wider rectangular patterns or circular patterns tended to generate cystic structures or multiple tubes (Fig. 2d).

In order to further analyse the effect of slit size and identify the suitable slit width, mathematical simulation was conducted. Three-dimensional cell deformation was simulated using a cell-centre model in which tissue shape was described by the positions of cell centres, with the epithelial sheet represented as a triangular network formed by neighbouring cell centres^23, 24^. Then, the single buckling was stably generated by cell proliferation when the slit width is smaller than 238.4 μm while it changes two-wave buckling mode when slit width larger (Fig. 2e). The estimated transition threshold from one-wave to two-wave mode was 260.5 μm (Fig. 2f).

Based on these design criteria, we constructed the final MoSAIC configuration using a modular polycarbonate frame containing a 300 µm-wide slit that narrowed to 150 µm at its opening (Fig. 2g). The cell-seeding area was enclosed by surrounding walls to maintain the defined tissue geometry. Matrigel was introduced into the slit and extended locally around its opening, where it was retained by surface tension and subsequently immobilized after gelation. hiPSCs were then seeded onto the upper surface of the platform and differentiated to generate spatially organized neural plate and surface ectoderm domains. The rigid polycarbonate frame enabled the entire cell–ECM construct to be manipulated intact with tweezers throughout culture and subsequently transferred into a fluidic chip for further spatial control, including establishment of the dorsal–ventral axis. Thus, MoSAIC integrates empirically and theoretically designed tissue geometry and ECM localization within a transferable modular architecture that enables additional spatial cues to be introduced sequentially during neural tube development.

### Developmental transition in neurulation

During neurulation *in vivo*, surrounding tissues provide spatially and temporally coordinated signals that pattern the developing neural tube. Sonic hedgehog (SHH) secreted from the notochord provides a ventralizing signal, whereas dorsal BMP signalling contributes to dorsoventral patterning^25^. In addition, studies in mice have shown that DKK1 is expressed during early neural tube closure and contributes to maintaining cells in an undifferentiated state during fusion and separation of the neural tube and surface ectoderm^21^. To reproduce these spatiotemporal changes *in vitro*, hiPSCs seeded on the polycarbonate device were first cultured in neural induction medium for 3 days. The device was then rotated by 90° and transferred into a fluidic chip, allowing the two sides of the tissue to be independently exposed to dorsal and ventral media. The dorsal medium contained BMP4 and a WNT inhibitor, whereas the ventral medium contained smoothened agonist (SAG), a pharmacological activator of SHH signalling, together with a BMP inhibitor and retinoic acid. This configuration established spatially distinct signalling environments across the developing tissue.

Figure 3a shows the transition of an initially continuous hiPSC sheet into two distinct tissues corresponding to neural tube and surface ectoderm. At day 1, cells predominantly expressed E-cadherin, whereas N-cadherin expression was barely detectable, and cells above the Matrigel-filled slit had begun to invaginate. By day 3, N-cadherin expression emerged within the invaginating population. By day 6, E-cadherin and N-cadherin became spatially segregated as invagination progressed, consistent with the epithelial remodelling associated with neurulation *in vivo*^26,27^. By day 10, the initially continuous sheet had bifurcated into two morphologically and molecularly distinct domains, with a tubular neural structure within the slit and a surrounding surface epithelial sheet. Figure 3b and Supplementary Video 3 show regions representing different stages of closure and tissue segregation. E-cadherin and cytokeratin 8 were predominantly restricted to the monolayer epithelial sheet on the platform, consistent with surface ectoderm identity, whereas the tubular structure within the slit expressed N-cadherin. In incompletely closed regions, the surface epithelial sheet remained continuous with the N-cadherin-positive neural tissue (Fig. 3b: section A–A’). In adjacent regions, the two tissues had fully segregated, forming a closed neural tube beneath a continuous surface ectoderm (Fig. 3b: section B–B’). Thus, intermediate and completed states of tissue bifurcation could be observed within the same construct.

**Figure 3.**
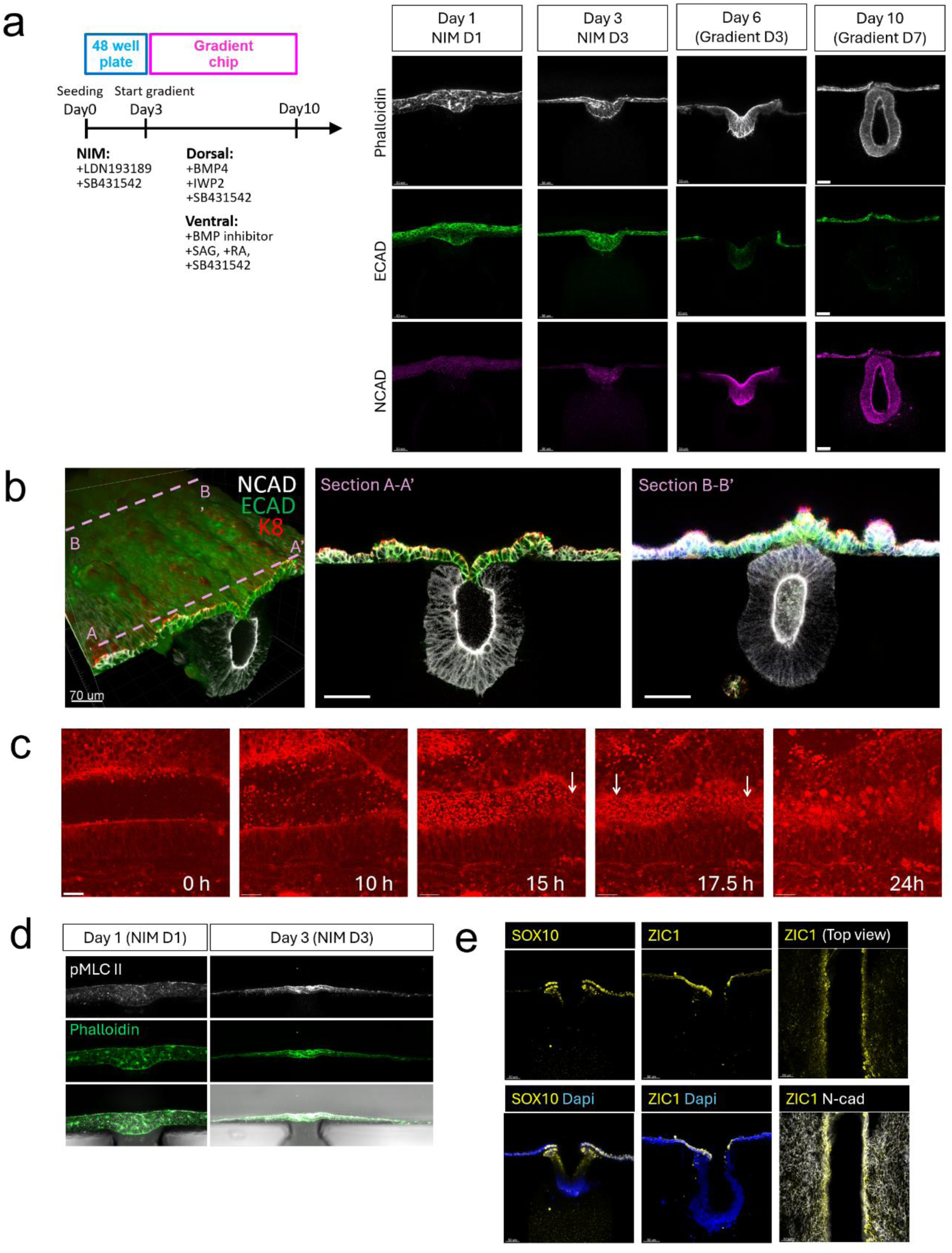
Developmental transition of a human iPSC sheet into a neural tube using MoSAIC. **a,** Time-course analysis of E-cadherin and N-cadherin expression during neural tube formation. E-cadherin was initially broadly expressed but gradually decreased in cells associated with the Matrigel-patterned region, whereas N-cadherin expression was initially weak and became evident in the Matrigel-associated region from day 6 onward. By day 10, the initially continuous epithelial sheet had segregated into two distinct tissues corresponding to the surface ectoderm and neural tube. Scale bar, 50 μm. **b,** Three-dimensional reconstruction of tissues stained for N-cadherin, E-cadherin and cytokeratin 8. Cross-section A–A′ shows a region with incomplete neural tube closure, whereas cross-section B–B′ shows a fully closed neural tube. Scale bar, 70 μm. **c,** Time-lapse imaging of actin dynamics during neural tube closure for 24 hours of culture. White arrows indicate actin-rich structures bridging the opposing tissue edges during closure. Scale bar, 30 μm. **d,** Spatial distribution of myosin II in the epithelial sheet during neural induction. Myosin II preferentially accumulated at the apical surface of cells within the Matrigel-associated region. **e,** Spatial distribution of SOX10 and ZIC1 before neural tube closure. Both markers were enriched at the boundary between the neural and non-neural ectodermal domains.

To directly visualize surface ectoderm closure, we developed a specialized gradient device compatible with time-lapse microscopy while maintaining separate dorsal and ventral culture environments (Supplementary Fig. 2). Two-photon imaging with SPY-actin (CY-SC202, Cytoskeleton Inc.) revealed progressive closure of an approximately 80 µm gap between opposing epithelial edges (Fig. 3c and Supplementary Video 4). After approximately 15 h, actin-rich protrusions bridged the gap at multiple positions, followed by contact and fusion of the opposing epithelial sheets. These observations enabled direct visualization of dynamic epithelial fusion during the human iPSC-derived neurulation process.

We next examined actomyosin organization during early invagination. At day 1, myosin II was distributed relatively uniformly, whereas by day 3 it became enriched in cells above the slit, particularly toward the apical side of the invaginating epithelium (Fig. 3d). Because apical actomyosin accumulation contributes to apical constriction and neural plate bending *in vivo*^20,28^, this enrichment coincided with the onset of tissue invagination in the MoSAIC model.

Neurulation also generates specialized cell populations at the boundary between neural and non-neural ectoderm^29–31^. Consistent with this process, SOX10, a neural crest marker, and ZIC1, associated with dorsal neural identities including the neural plate border and roof plate, were detected at the boundary region (Fig. 3e), indicating acquisition of molecular features characteristic of the neural– non-neural ectoderm interface.

Collectively, the MoSAIC-derived model recapitulated a sequence of developmental transitions associated with neurulation, including actomyosin enrichment, tissue invagination, cadherin switching, emergence of neural border-associated identities, epithelial fusion, and bifurcation of a continuous ectodermal sheet into neural tube and surface ectoderm. Rather than generating these tissues independently as terminal structures, MoSAIC reconstructed their emergence through a continuous morphogenetic transition.

### Spatial expression pattern

To evaluate the spatial organization of the MoSAIC-derived neural tube, we examined anterior– posterior (AP) identity, apical–basal polarity, and dorsoventral (DV) patterning after 10 days of culture by immunostaining.

Although no AP morphogen gradient was imposed, regional identity could emerge from the differentiation conditions. We therefore examined OTX2, HOXB1, and HOXC9 expression to identify the axial identity of the resulting tissue (Fig. 4a). OTX2, associated with anterior neural identity, was undetectable, whereas HOXC9 was strongly expressed in the neural tube, with weak HOXB1 expression. This profile suggests a predominantly posterior spinal cord-like identity rather than an anterior neural or hindbrain identity. We next assessed apical–basal polarity by staining for ZO-1 and INTEGRIN β1 (Fig. 4b). ZO-1 was concentrated along the luminal surface, whereas INTEGRIN β1 localized predominantly to the outer basal surface of the neural tube. This complementary distribution confirmed the establishment of apical–basal polarity, with the apical surface facing the lumen and the basal surface contacting the surrounding ECM.

**Figure 4.**
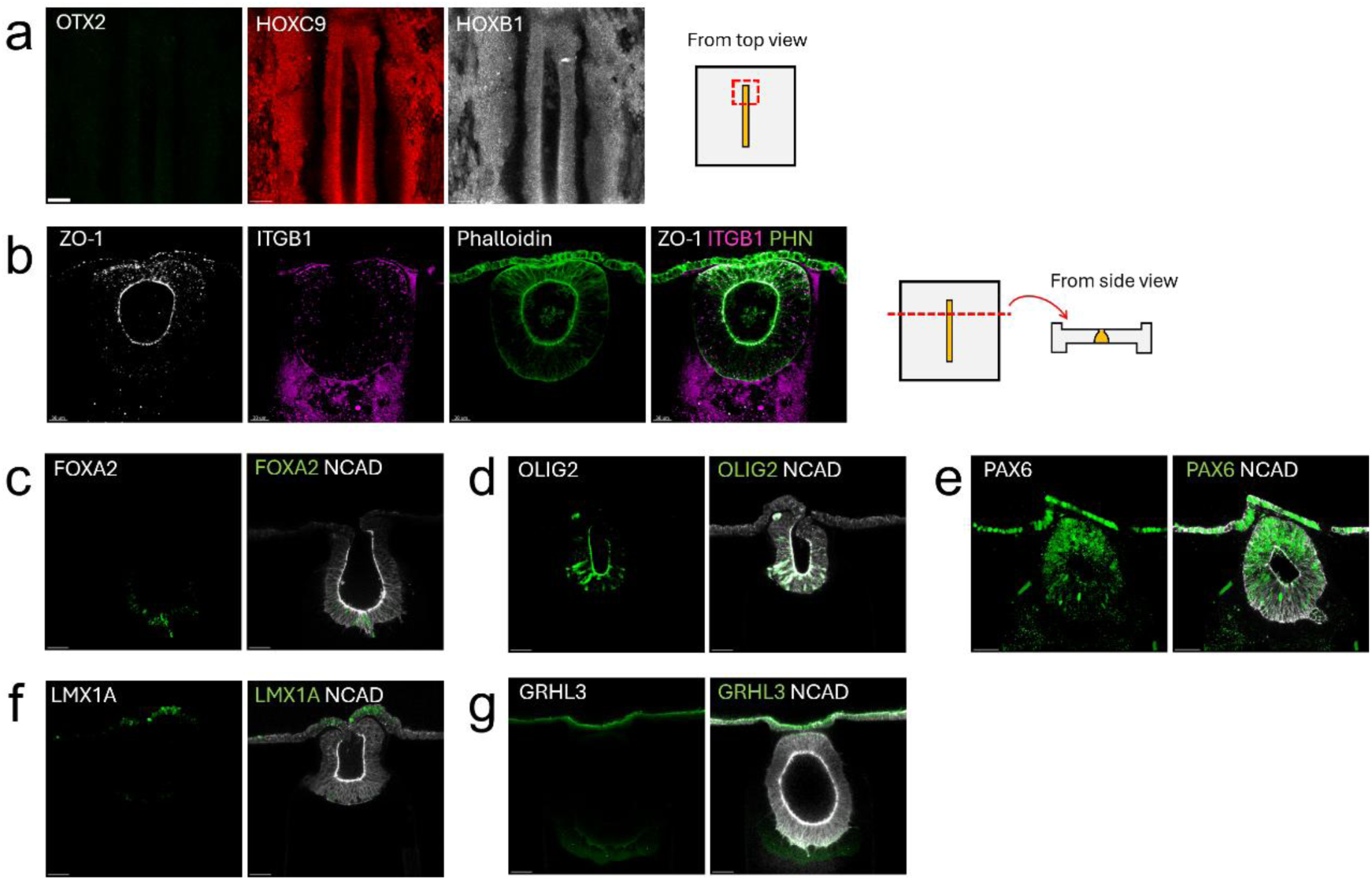
Spatial patterning and epithelial polarity of neural tubes generated using MoSAIC. **a,** Immunofluorescence staining for OTX2, HOXB1 and HOXC9 to assess the anterior–posterior (AP) identity of the generated neural tubes. **b,** Assessment of apical–basal polarity by immunofluorescence staining for ZO-1 and integrin β1, together with phalloidin staining to visualize filamentous actin (F-actin). **c–g,** Immunofluorescence staining to assess dorsal–ventral (DV) patterning, showing the spatial distributions of FOXA2 (c), OLIG2 (d), PAX6 (e), LMX1A (f) and GRHL3 (g). Scale bar, 50 μm.

During neurulation *in vivo*, opposing morphogen signals establish spatially distinct neural progenitor domains along the DV axis^32, 33^. We therefore examined the distribution of key DV markers (Fig. 4c–g). In the developing neural tube, FOXA2 marks the ventral floor plate, OLIG2 identifies adjacent motor neuron progenitors, and PAX6 is expressed in intermediate and dorsal progenitor domains. The MoSAIC-derived neural tube reproduced this relative organization, with FOXA2-positive cells concentrated ventrally, OLIG2-positive cells positioned adjacent to them, and PAX6-positive cells distributed further dorsally, which were consistent with DV progenitor patterning observed *in vivo*. LMX1A-positive cells were also detected dorsally but were predominantly located within the surface ectoderm rather than the dorsal neural tube (Fig. 4c–g). Thus, a distinct LMX1A-positive roof-plate domain was not established under the present conditions. GRHL3 expression was similarly detected in dorsal and surface ectodermal regions, consistent with its involvement in dorsal neural tube and non-neural ectodermal programs.

Collectively, these results demonstrate that MoSAIC-derived neural tissue acquired posterior spinal cord-like identity, apical–basal polarity, and spatially ordered DV progenitor domains. The relative organization of FOXA2, OLIG2, and PAX6 indicates that engineered morphogen environments established positional information within the neural tube, although the expanded ectodermal localization of LMX1A highlight remaining differences from *in vivo* patterning.

### Mechano-chemical perturbation to analyse the neural tube closure process

A major advantage of *in vitro* human models is the ability to independently manipulate environmental cues that are difficult to perturb *in vivo*. We therefore investigated how mechanical and chemical spatial cues influence neural tube morphogenesis using MoSAIC.

We first examined the contribution of ventral Sonic hedgehog (SHH) signalling to neural tube morphogenesis. Previous studies in mouse and chick embryos have shown that notochord removal does not necessarily prevent neural tube closure, although notochord-derived SHH is essential for ventral neural patterning and contributes to tissue morphogenesis^34-36^. To investigate this relationship in a human model, we manipulated the spatial distribution of smoothened agonist (SAG), a SHH pathway activator (Fig. 5a, b). Under the standard dorsoventral (DV) gradient with ventral SAG, the neural tube exhibited prominent ventral extension, forming an elongated morphology (Fig. 5a(i)). Removing SAG from the ventral medium significantly reduced the DV length, producing a more rounded structure (Fig. 5a(ii), b). Nevertheless, neural tube closure occurred both without SAG and under dorsal-medium-only conditions (Fig. 5a(ii, iii)), indicating that ventral SHH pathway activation was not strictly required for closure in this model. In contrast, introducing SAG into the dorsal medium markedly disrupted tissue morphogenesis (Fig. 5a(iv–vi)). Under these conditions, most samples exhibited little or no invagination into the slit, instead maintaining a relatively flat epithelial sheet resembling the morphology observed immediately after neural induction at day 3. Notably, dorsal SAG exposure did not induce substantial tissue bending toward the dorsal side. These observations suggest that SHH signalling does not independently determine the direction of tissue bending but acts in conjunction with the surrounding DV signalling environment to regulate tissue deformation and ventral extension.

**Figure 5.**
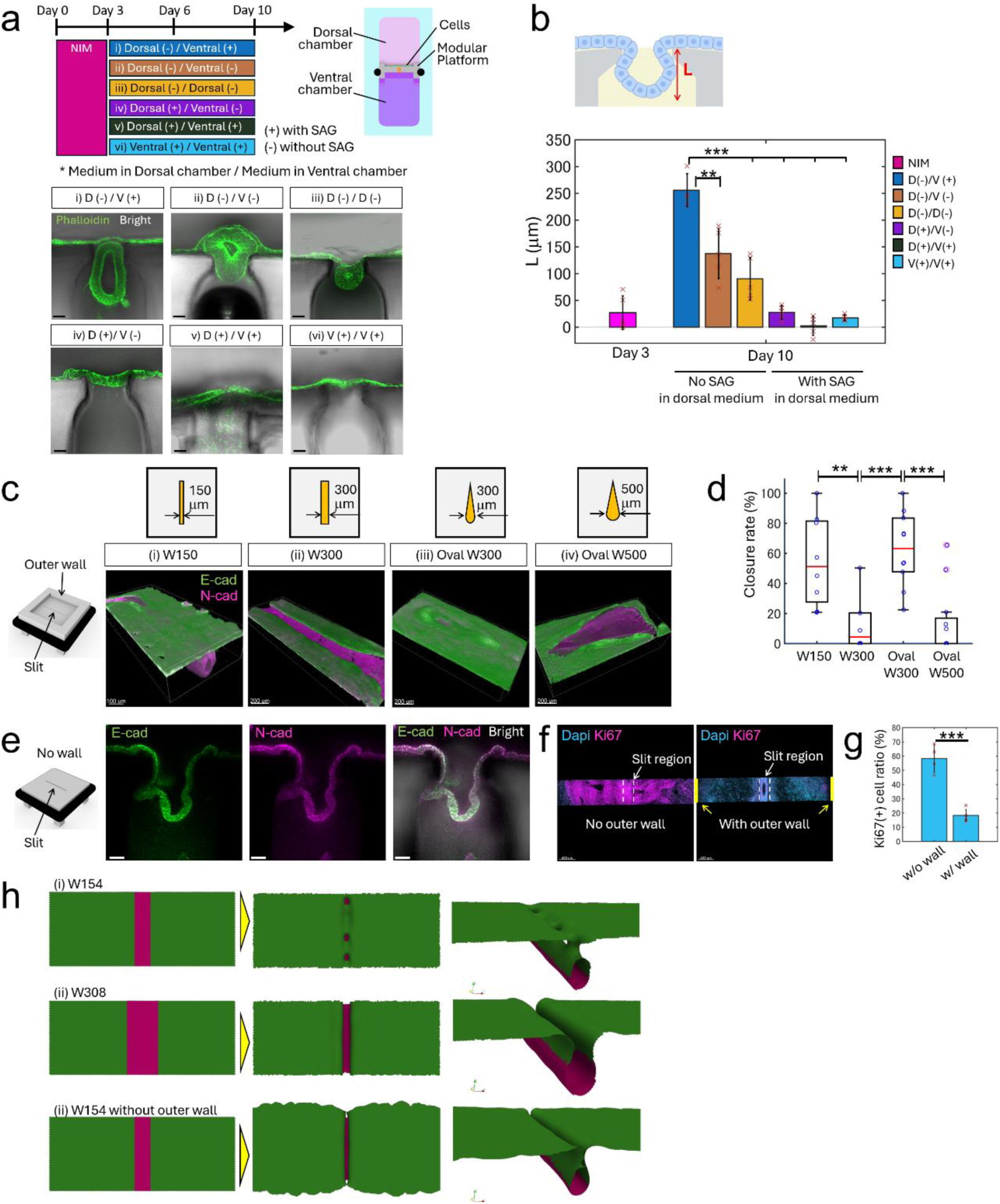
Mechanochemical perturbation of neural tube morphogenesis using MoSAIC. **a,** Chemical perturbation to investigate the contribution of spatially controlled SHH signalling to neural tube morphogenesis. Six culture conditions were compared: (i) dorsal (−)/ventral (+), (ii) dorsal (−)/ventral (−), (iii) dorsal (−)/dorsal (−), (iv) dorsal (+)/ventral (−), (v) dorsal (+)/ventral (+) and (vi) ventral (+)/ventral (+). The (+) and (−) symbols indicate the presence and absence of the SHH pathway agonist SAG, respectively, while the terms before and after the slash (/) indicate the media supplied to the dorsal and ventral chambers, respectively. Scale bar, 50 μm. **b,** Quantification of neural tube elongation along the dorsal–ventral (DV) axis under different SAG exposure conditions (n=5). Ventral-specific SAG exposure significantly enhanced DV elongation compared with conditions lacking spatially localized SAG signalling. **c,** Representative neural tube morphologies generated using slits with different widths and geometries. Slit dimensions and geometry defined the initial neural plate region and influenced subsequent neural tube closure. **d,** Quantification of neural tube closure efficiency as a function of slit width and geometry (n>5). **e,** Representative cross-sectional images of tissues cultured using a modular device without the outer wall. Although tissue invagination into the Matrigel-filled slit occurred, segregation into distinct neural tube and surface ectoderm tissues was impaired. Scale bar: 50 μm. **f,** Immunofluorescence staining for Ki67 on day 6 of culture using modular devices with and without the outer wall. **g,** Quantification of the proportion of Ki67-positive cells relative to the total cell population in the outer quarter of the culture region, comparing devices with and without the outer wall (n>4). **h,** Mathematical simulations of tissue morphogenesis under different geometric constraints: (i) a 150-μm-wide slit with the outer wall, (ii) a 300-μm-wide slit with the outer wall and (iii) a 150-μm-wide slit without the outer wall. The simulations incorporated contact inhibition of cell proliferation and reproduced the geometry-dependent morphological differences observed experimentally. **P < 0.01; ***P < 0.001.

We next examined the effects of physical boundary. Figure 5c shows the morphological results with various slit dimensions and shapes after 10 days of culture. With a 150 µm-wide rectangular slit, approximately 50% of the neural tube length was closed on average, with complete closure observed in some samples (Fig. 5c(i)). In contrast, a 300 µm-wide rectangular slit frequently resulted in incomplete or minimal closure (Fig. 5c(ii)). Interestingly, an oval slit with a maximum width of 300 µm exhibited closure efficiency comparable to that of the 150 µm-wide rectangular slit (Fig. 5c(iii)), whereas increasing the oval width to 500 µm markedly impaired closure (Fig. 5c(iv)). Quantification confirmed that closure efficiency depended on both the dimensions and geometry of the patterned neural region (Fig. 5d).

Beyond the slit geometry itself, we found that the outer boundary of the modular platform also provided an important physical constraint. When the surrounding outer wall was removed, the cell sheet deformed at multiple locations across the platform and failed to undergo spatially confined segregation into a single tubular structure (Fig. 5e). This indicates that the outer wall contributes to the mechanical boundary conditions required to direct tissue deformation toward the Matrigel-filled slit rather than allowing unconstrained expansion of the epithelial sheet. To further examine how the outer boundary influenced cell behaviour, we quantified Ki67-positive cells after 6 days of culture on platforms with or without the surrounding wall (Fig. 5f). The proportion of Ki67-positive cells relative to the total cell population was significantly lower in constructs confined by the outer wall than in those cultured without the wall (Fig. 5g). This reduction in proliferation is consistent with increased contact inhibition under spatial confinement, whereas cells on platforms without an outer wall could expand outward and thereby relieve local crowding. These results suggest that the outer boundary not only defines tissue geometry but also alters the proliferative state and mechanical environment of the epithelial sheet.

Together, these perturbation experiments demonstrate that neural tube closure and morphology can be differentially regulated. Closure efficiency was strongly influenced by neural plate geometry and physical confinement, whereas spatial SHH pathway activation modulated DV extension and invagination, with ectopic dorsal activation disrupting tube formation. These findings establish MoSAIC as a platform for systematically dissecting the contributions of geometric constraints and spatial biochemical signalling to human neural tube morphogenesis.

The quantitatively defined geometry of MoSAIC further enabled direct integration with mathematical modelling. Simulations incorporating cell proliferation and density-dependent contact inhibition were performed under three experimentally matched boundary conditions: a 154 µm slit, a 308 µm slit, and a 154 µm slit without the outer wall (Fig. 5h). The simulations reproduced the characteristic deformation patterns observed experimentally under each condition, supporting the relationship between tissue geometry, external confinement, and morphogenesis. Together with the reduced Ki67-positive fraction observed under confined conditions, these results demonstrate that experimentally defined boundary conditions in MoSAIC can be directly translated into computational models, providing a basis for future quantitative and predictive analyses of developmental morphogenesis.

## Discussion

Development proceeds through coordinated transitions in which tissues change shape, acquire positional information, segregate into distinct domains, and establish new interfaces. Reconstructing these processes *in vitro* remains challenging because developmental microenvironments integrate multiple spatial cues that act simultaneously and change over time. Here, we developed MoSAIC, a modular platform that combines rigid polycarbonate with soft ECM hydrogels to independently organize tissue geometry, ECM localization, physical boundaries, and soluble-factor distribution. By decomposing complex culture operations into controllable modules and subsequently integrating them, MoSAIC provides a framework for reconstructing dynamic developmental environments rather than only specifying endpoint tissue identities.

Using MoSAIC, we reconstructed a continuous transition from an initially uniform hiPSC-derived epithelial sheet into neural tube and surface ectoderm. The model reproduced sequential features of neurulation, including neural specification, invagination, cadherin switching, epithelial fusion, tissue bifurcation, apical–basal polarity, and dorsoventral patterning. Intermediate states of closure were directly observed, and live imaging captured progressive fusion of the surface ectoderm, enabling the developmental transition itself to be experimentally accessed.

MoSAIC also enables controlled perturbation of individual environmental inputs. Altering neural plate geometry and physical confinement strongly affected closure efficiency, whereas removal of ventral SAG did not abolish closure but altered dorsoventral extension and tube morphology. These results show that neural tube closure and overall tube morphology are regulated by partly distinct environmental factors, highlighting the utility of MoSAIC for independently analysing mechanical and biochemical contributions to morphogenesis.

The present model nevertheless captures only part of human neurulation. The neural tube retained a relatively simple elliptical cross-section rather than the more expanded morphology observed *in vivo*. Although posterior neural identity and dorsoventral progenitor organization were established, it remains unclear whether the tissue can subsequently generate and spatially organize differentiated neuronal populations such as motor and sensory neurons. Further optimization of axial patterning^37–39^, planar cell polarity^40–42^, mechanical constraints, and long-term culture will therefore be required to reproduce later neural tube maturation.

More broadly, MoSAIC shifts organoid engineering from endpoint specification toward process-oriented developmental modelling. By enabling spatial cues to be assembled, visualized, modified, and perturbed within a single platform, it provides a framework for investigating how geometry, ECM organization, and morphogen signalling cooperate to drive human morphogenesis.

## Methods

### hiPSC culture

IMR90-4 iPSCs (WiCell, WB65317) were maintained on Matrigel-coated dishes in mTeSR Plus medium (STEMCELL Technologies, 100-0276) and dissociated using ReLeSR (STEMCELL Technologies, 05872). For Matrigel coating, growth factor reduced phenol red free (Corning, 356231) was diluted 1:100 in DMEM/F12, and 1.5 mL of the diluted Matrigel solution was added to each well of a 6-well plate. The plate was incubated at room temperature for 1 h before use.

### 2D cell and ECM control for E-cadherin / N-cadherin localization

Photolithography was employed to produce a shadow mask membrane in order to pattern cells on a dish. A silicon substrate was ultrasonicated with isopropyl alcohol and deionized water individually to clean the substrate. A SUEX100 (DJ Microlaminates, MA, U.S.A) negative thick dry photoresist was used to lithograph the designed pattern to the substrate. Once the pattern has been transferred to the photoresist, the silicon substrate was used as a mould for producing a PDMS membrane with the pattern. PDMS (Silpot 184, Dow Toray, 04133124) was poured onto the mould at a thickness lower than that of the mould and incubated in the oven at 90°C for 20 min. After the PDMS membrane had been cured, it was detached from the silicon mould, and 70% ethanol was sprayed on the membrane and a culture dish. Subsequently, cell suspension medium was applied over the membrane in the unmasked area, and the dish was incubated at 37°C for 2 hours. The PDMS membrane was then removed for further experiments (Supplementary Fig. 4). Cells were cultured with neural induction medium for 3 days followed by 2 days of additional culture with dorsal medium.

### Quantifications of E-cadherin/N-cadherin positive region

Regions expressing E-cadherin or N-cadherin were quantified from fluorescence images. Because both E-cadherin and N-cadherin are localized at the cell membrane, their fluorescence signals appeared as thin outlines along cell borders. Therefore, cell-based segmentation was not performed. Instead, images were segmented based on expression patterns using ImageJ.

To distinguish regions with different expression patterns, background subtraction was first performed using a rolling ball radius of 50 pixels. A Gaussian blur (σ = 10 µm) was then applied to smooth the fluorescence signals and enable extraction of continuous expression domains from the mosaic-like staining pattern (Supplementary Fig. 5).

After image preprocessing, the entire processed image was selected for analysis, and a plot profile was generated. From the resulting intensity profile, fluorescence intensities were sampled at 50 µm intervals from the centre of the slit. The extracted data were then imported into MATLAB to generate fluorescence intensity distribution graphs.

### MoSAIC culture for neurulation

A schematic overview of the MoSAIC culture procedure is provided in Supplementary Fig. 3. The modular device was designed using 3D CAD software (Rhinoceros) and fabricated from polycarbonate by machining. A silicone gasket was attached to the device to ensure sealing upon integration with the fluidic chip. Before cell seeding, the backside of the device was plasma-treated, and 0.5 µl of Matrigel was introduced into the slit region. The device was incubated at 37 °C for 3 min to allow gelation. Subsequently, 20 µl of 4% Matrigel diluted in DMEM was applied to the top surface of the device and incubated at room temperature for 1 h to coat the cell-seeding surface.

After removal of the Matrigel-containing DMEM, 20 µl of cell suspension in mTeSR medium was applied to the device at a density of 2.0–6.0 × 10^6^ cells/ml. Cells were incubated at 37 °C for 2 h to allow attachment. Excess medium was then removed, and the device was transferred to a 48-well plate containing neural induction medium and cultured for 3 days.

After the initial 3-day culture period, the cell-laden device was transferred to the fluidic chip. The device was placed in the chip base with the silicone gasket positioned in the corresponding groove. The chip lid and base were sealed using a PDMS double-sided adhesive film (NSD-100, NIPPA), in which openings were created to provide access to the medium chambers. After assembly, dorsal medium was introduced into the chamber on the top side of the device, whereas ventral medium was introduced into the chamber on the opposite side. This configuration enabled independent control of the two medium compartments and established spatially defined dorsal and ventral culture conditions across the developing tissue.

### Differentiation protocols

For neural induction and subsequent dorsoventral patterning, a basal medium consisting of 49% Neurobasal Plus medium (Invitrogen, A3582901), 49% KnockOut DMEM/F-12 (Invitrogen, 12660012), 1% GlutaMAX Supplement (Invitrogen, 35050061), and 1% MEM Non-Essential Amino Acids Solution (100×; Invitrogen, 11140050) was used.

For neural induction, hiPSCs adhered to the modular device were cultured in a 48-well plate for 3 days in basal medium supplemented with 2 µM LDN193189 (R&D Systems, 5631/10) and 2 µM SB431542 (Nacalai Tesque, 18176-54). Following neural induction, the device was transferred to a fluidic chip for an additional 7 days to establish dorsoventral patterning. Dorsal medium containing 1 ng/mL BMP4 (PeproTech, 120-05), 3 µM IWP2 (R&D Systems, 3533/10), and 2 µM SB431542 was supplied to the dorsal chamber, whereas ventral medium containing 1 µM LDN193189, 300 nM all-trans retinoic acid (RA; Nacalai Tesque, 36331-31), 500 nM smoothened agonist (SAG; Enzo Life Sciences, ALX-270-426-M001), and 2 µM SB431542 was supplied to the ventral chamber. Both dorsal and ventral media were prepared using the same basal medium. Throughout the 7-day gradient culture, the media in both chambers were replaced daily.

### Immunostaining and Imaging

Samples were washed with Dulbecco’s phosphate-buffered saline (DPBS) and fixed with 4% paraformaldehyde for 10 min, followed by two washes with DPBS for 10 min each. For immunofluorescence staining, samples were permeabilized with 0.5% Triton X-100 for 20 min and subsequently washed three times with 100 mM glycine for 10 min each. The immunofluorescence (IF) buffer consisted of 0.05% Tween-20, 0.2% Triton X-100, and 0.1% bovine serum albumin (BSA; Sigma, 126615) in DPBS. Samples were blocked for 50 min in IF buffer supplemented with 1% donkey serum (Merck, D9663), hereafter referred to as IF+G buffer.

Primary antibodies were diluted in IF+G buffer at the concentrations listed in Table 1 and incubated with the samples overnight at 4 °C. Secondary antibodies were similarly diluted in IF+G buffer and incubated for 2 h at room temperature. Following each antibody incubation, samples were washed three times with IF buffer for 20 min each. Nuclei were stained with DAPI for 2 h, followed by three washes with DPBS for 5 min each. For three-dimensional (3D) imaging, samples were optically cleared overnight using RapiClear clearing solution (SUNJin Lab, RC149001) before imaging.

Fluorescence imaging of both sectioned and intact 3D samples was performed using a Leica TCS SP8 confocal microscope equipped with a 25× water-immersion objective (HC FLUOTAR L 25×/0.95 W VISIR). Time-lapse imaging was performed using an EVIDENT FV-3000, FVRS-F2SJ multiphoton microscope equipped with a 25× water-immersion objective (XLPLN25XWMP: NA1.05).

### Cryosectioning

For selected samples, cryosectioning was performed before immunostaining. Following fixation, samples were immersed in 10% sucrose (Wako, 194-00011) for 1 h, followed by 30% sucrose overnight, while remaining attached to the modular polycarbonate platform. Samples were then incubated in optimal cutting temperature (O.C.T.) compound (Tissue-Tek; Sakura Finetek, 4583) for 10 min before embedding in fresh O.C.T. compound and freezing at −80 °C for 30 min. The frozen samples, including the modular polycarbonate platforms, were sectioned at a thickness of 100–200 µm using a cryostat (Leica CM3050 S) equipped with a specimen freezing unit. To prevent tissue detachment from the polycarbonate platform, 1.5% agarose solution was applied to the sections collected on glass slides. After agarose gelation, the sections were washed with Milli-Q water to remove residual O.C.T. compound.

### Mathematical simulation

Mathematical simulations were performed for cell invagination and tube formation to examine how morphology depended on slit width and boundary conditions. The detailed conditions and explanation are described in Supplementary Information.

### Statistical analysis

Sample numbers for a given experiment are provided in each figure legend and were always *n* ≥ 4 independent replicates. Statistical significance between two independent groups was assessed using a two-tailed Welch’s t-test without assuming equal variances. Statistical analysis was performed using Matlab.

## Acknowledgements

This research was supported by funds from JSPS KAKENHI grants 23KK0205, 24H00806, 26H01002 and AMED under Grant Number JP26bm1123073. The multi-photon time-lapse imaging experiment was performed at the Kobe BioImaging Facility and Factory.

## Contributions

M.H conceptualized MoSAIC system and designed for neurulation. M.H and I.M designed culture conditions for neural tube development from iPSCs. M.H, S.N, Y.S, performed experiments for neural tube developments and collecting data to validate the transition process and functionality at the endpoint of neural tube development. L.Y.G.T and I.K conducted cell culture to backup the simulation results and design slit width. Y.X.N conceived sectioning of composite material of ECM and polycarbonate and its analysis. T.M and Y.I developed mathematical models to design the environment and analyse the effect of environmental perturbations. M.H wrote the manuscript with review and feedback from all other authors.

---

**Supplementary Figure 1.**
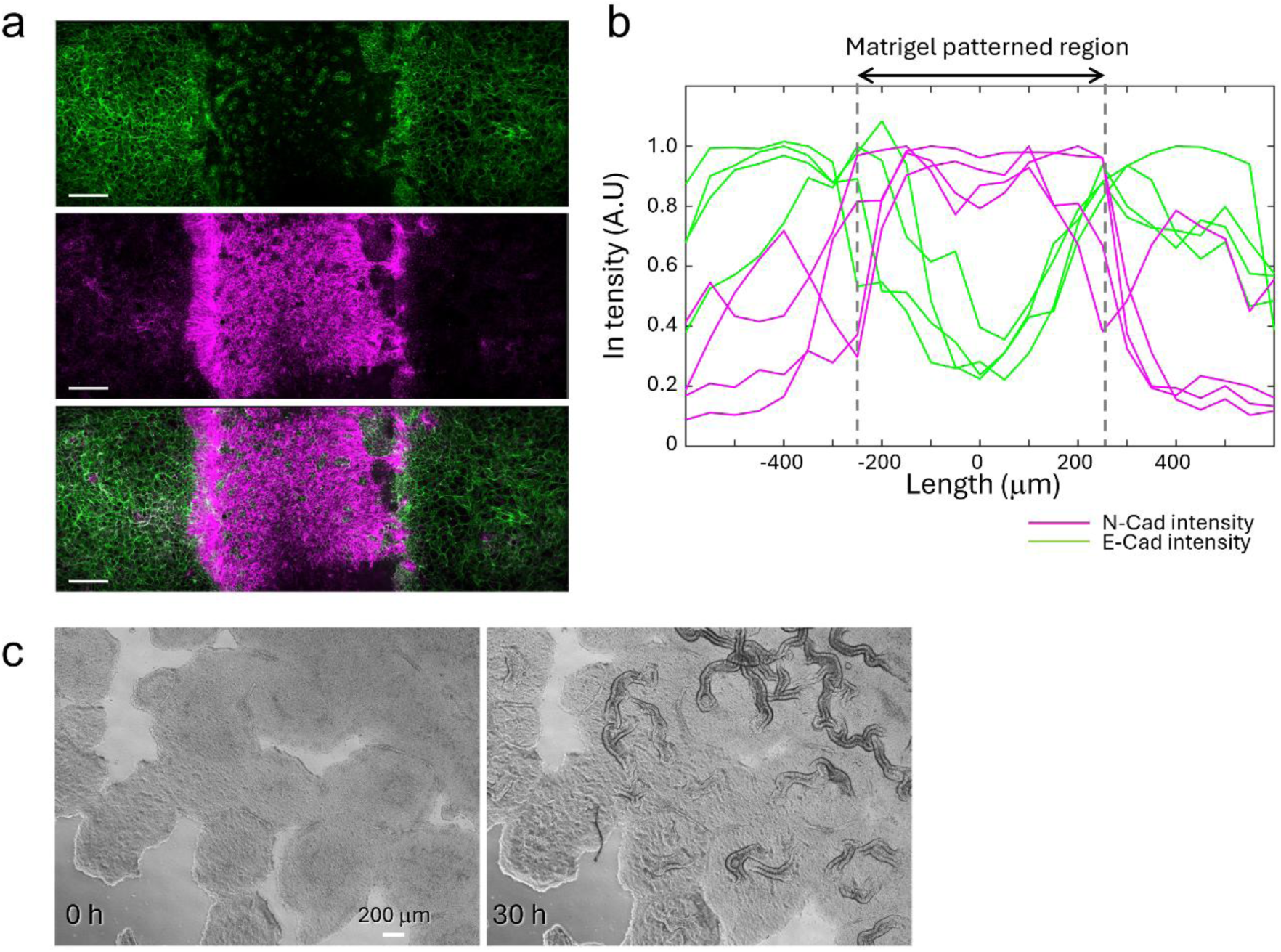
Spatial regulation of neural differentiation by ECM patterning and spontaneous wrinkle formation during neural induction. **a,** Immunofluorescence images showing the spatial distributions of E-cadherin and N-cadherin in hiPSCs cultured with a 500-μm-wide rectangular Matrigel pattern. N-cadherin-positive cells emerged within the Matrigel-patterned region, whereas cells outside this region predominantly expressed E-cadherin. Compared with the 250-μm-wide Matrigel pattern (Fig. 2a), the N-cadherin-positive region exhibited a broader distribution. Scale bar, 100 μm. **b,** Quantification of the spatial distributions of E-cadherin- and N-cadherin-positive regions. The dot lines indicate the boundary of the Matrigel region. **c,** Time-lapse imaging of hiPSCs cultured on a conventional dish under neural induction conditions. Multiple wrinkle patterns spontaneously emerged across the epithelial sheet during neural induction (Supplementary Video 2).

**Supplementary Figure 2.**
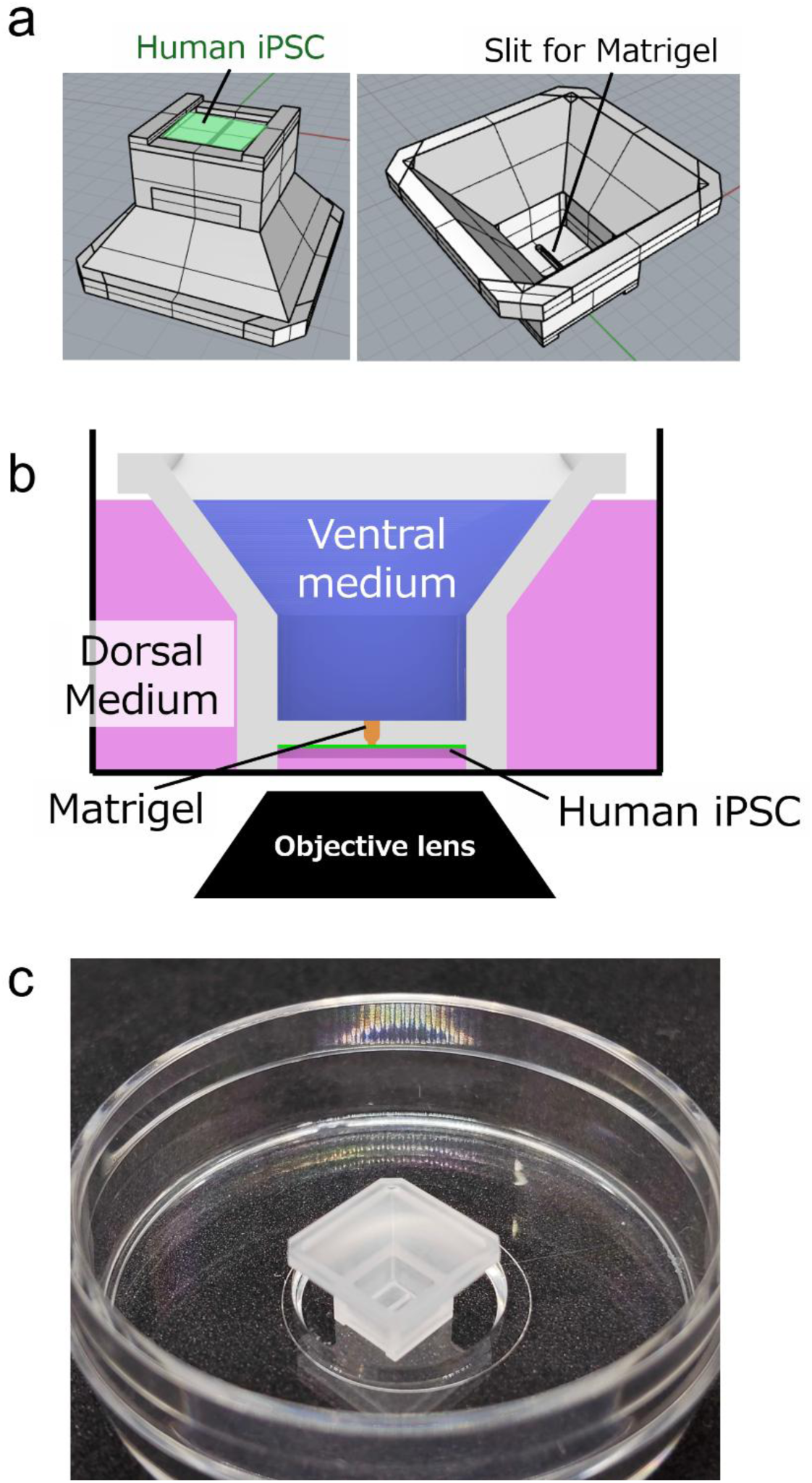
Design of a specialized gradient device for time-lapse imaging of neural tube morphogenesis. **a,** Schematic illustration of the device design by CAD. A slit with the same geometry used for neural tube formation (150 μm wide at the entrance and 300 μm wide at the exit) was incorporated into the device, with cells seeded on its bottom surface. b, Schematic illustration of the culture configuration. Matrigel was localized within the slit, while dorsal and ventral media were spatially compartmentalized by the device. The cell-seeded surface was oriented toward the objective lens to enable time-lapse imaging of neural tube morphogenesis. c, Photograph of the assembled device placed in a glass-bottom dish.

**Supplementary Figure 3.**
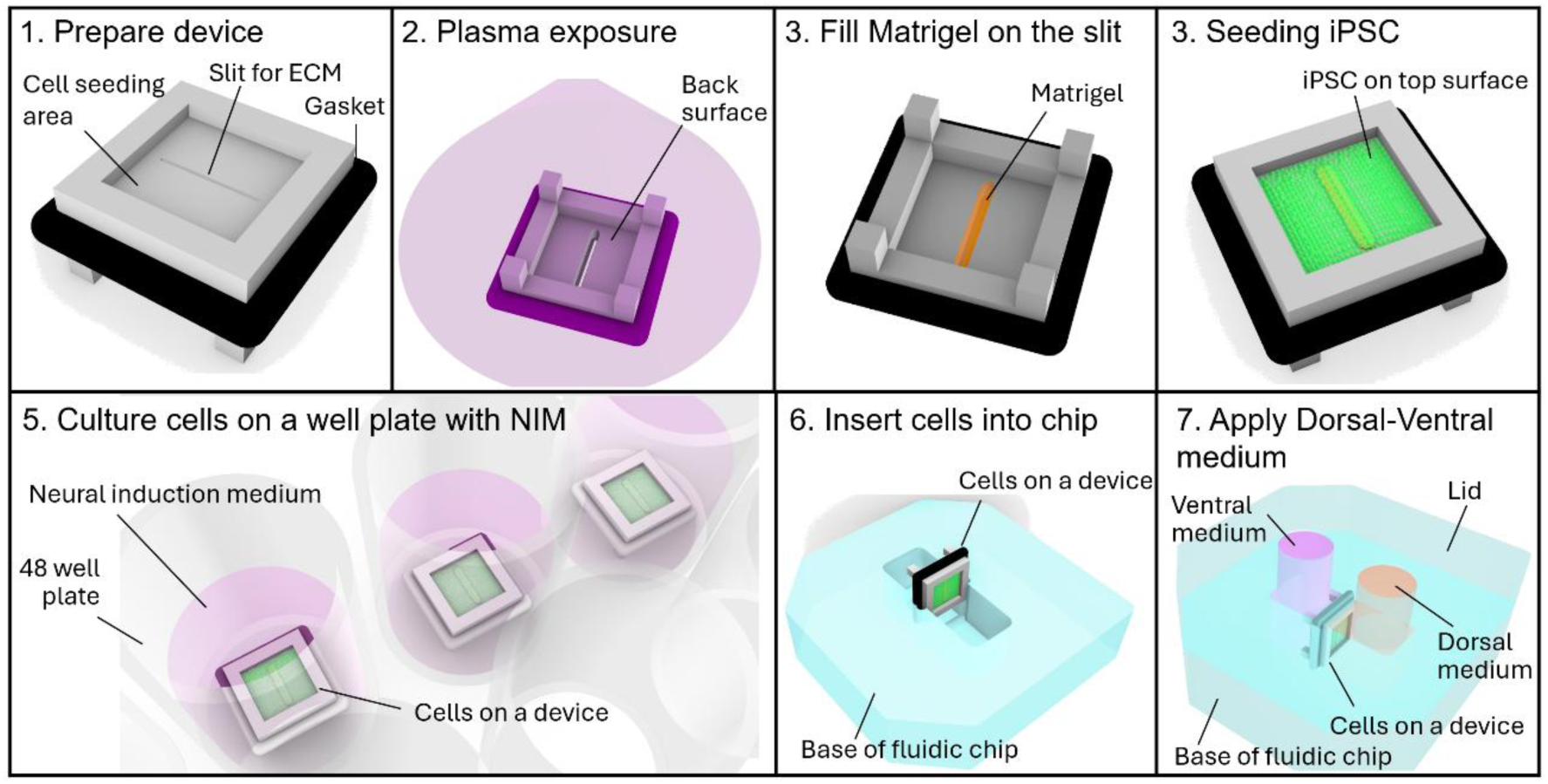
Schematic illustration of the MoSAIC workflow for integrating multimodal spatial cues.

**Supplementary Figure 4.**
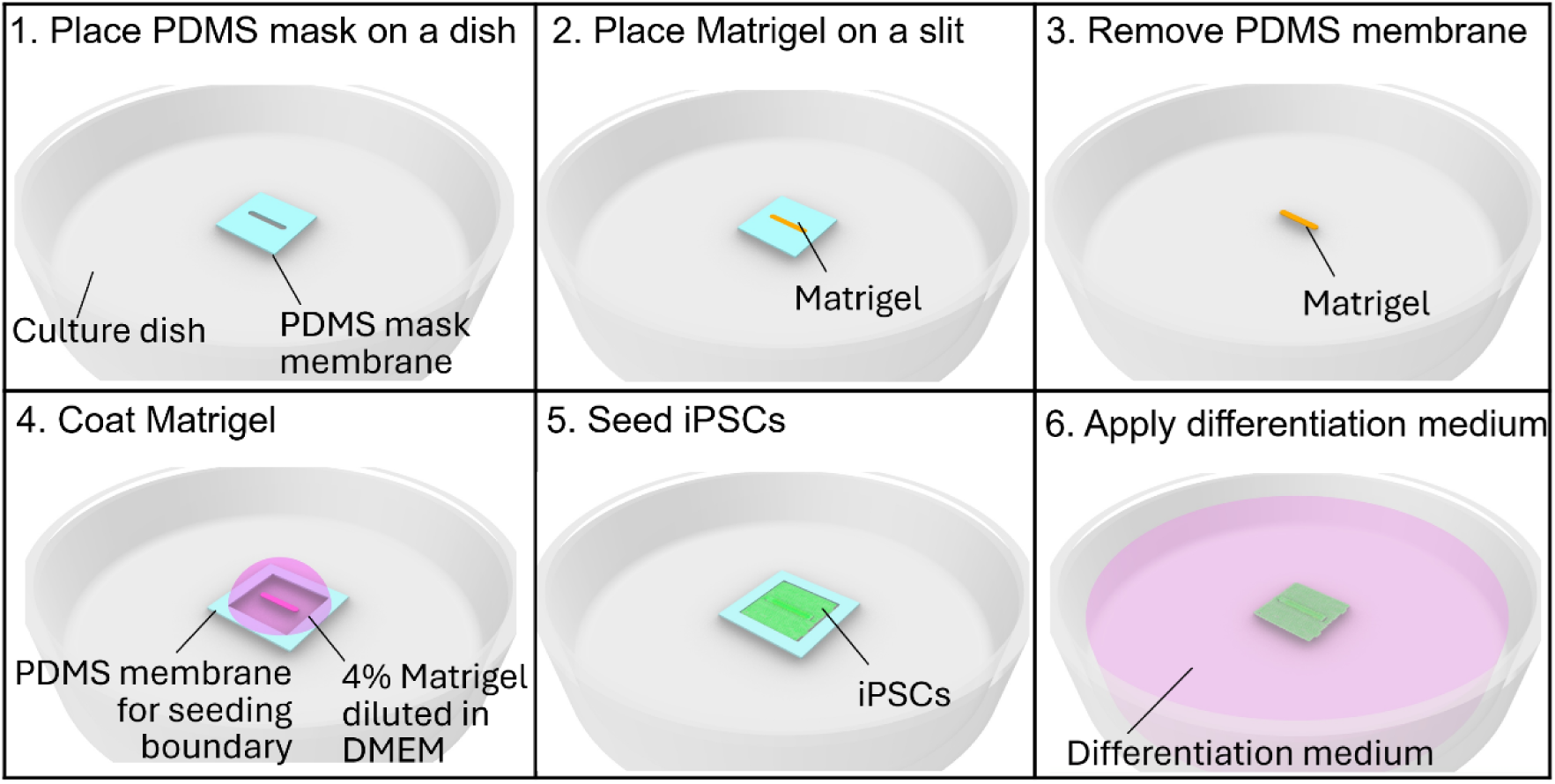
Schematic illustration of the ECM localization procedure on a glass-bottom dish.

**Supplementary Figure 5.**
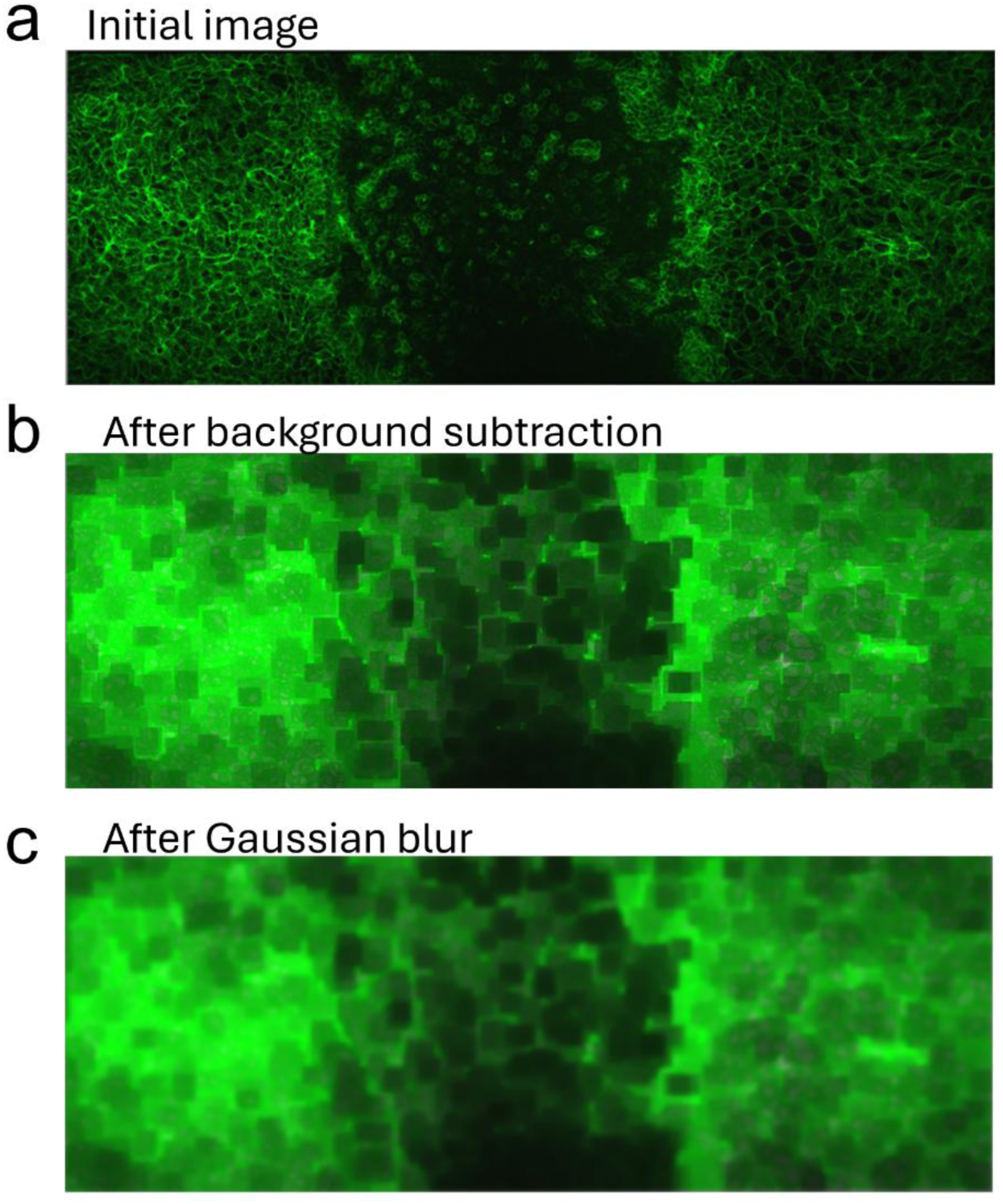
Image-processing workflow for quantification of E-cadherin- and N-cadherin-positive regions. **a,** Original fluorescence image. **b,** Image after background subtraction using a rolling-ball radius of 50 pixels. **c,** Image after Gaussian smoothing (σ = 10 µm).

## References

1. Shahbazi, M. N. Mechanisms of human embryo development: from cell fate to tissue shape and back. Development 147, dev190629 (2020).

2. Zhu, M. & Zernicka-Goetz, M. Principles of self-organization of the mammalian embryo. Cell 183, 1467–1478 (2020).

3. Nikolopoulou, E., Galea, G. L., Rolo, A., Greene, N. D. E. & Copp, A. J. Neural tube closure: cellular, molecular and biomechanical mechanisms. Development 144, 552–566 (2017).

4. Di Lullo, E. & Kriegstein, A. R. The use of brain organoids to investigate neural development and disease. Nat. Rev. Neurosci. 18, 573–584 (2017).

5. Sun, Y. et al. Bioengineering innovations for neural organoids with enhanced fidelity and function. Cell Stem Cell 32, 689–709 (2025).

6. Andersson-Rolf, A. & Clevers, H. New developments and applications of human organoids. Nat. Rev. Mol. Cell Biol. 27, 543–558 (2026).

7. Hofer, M. & Lutolf, M. P. Engineering organoids. Nat. Rev. Mater. 6, 402–420 (2021).

8. Karzbrun, E. et al. Human neural tube morphogenesis *in vitro* by geometric constraints. Nature 599, 268–272 (2021).

9. Xue, X. et al. A patterned human neural tube model using microfluidic gradients. Nature 628, 391–399 (2024).

10. Pagliaro, A. et al. Temporal morphogen gradient-driven neural induction shapes single expanded neuroepithelium brain organoids with enhanced cortical identity. Nat. Commun. 14, 7361 (2023).

11. Nikolaev, M. et al. Homeostatic mini-intestines through scaffold-guided organoid morphogenesis. Nature 585, 574–578 (2020).

12. Onesto, M. M., Kim, J.-I. & Pașca, S. P. Assembloid models of cell–cell interaction to study tissue and disease biology. Cell Stem Cell 31, 1563–1573 (2024).

13. Kim, J.-I. et al. Human assembloid model of the ascending neural sensory pathway. Nature 642, 143–153 (2025).

14. Reza, H. A. et al. Multi-zonal liver organoids from human pluripotent stem cells. Nature 641, 1258–1267 (2025).

15. Graham, A. J. et al. Stress-relaxing granular bioprinting materials enable complex and uniform organoid self-organization. Nat. Mater. 25, 1239–1251 (2026).

16. Wilt, J. K., Larson, N. M. & Lewis, J. A. Rotational multimaterial 3D printing of soft robotic matter with embedded asymmetrical pneumatics. Adv. Mater. 38, e10141 (2026).

17. Jowett, G. M. et al. ILC1 drive intestinal epithelial and matrix remodelling. Nat. Mater. 20, 250–259 (2021).

18. Guo, J. et al. 3D biomimetic niche modulates embryo development *in vitro*. Nat. Commun. 17, 1279 (2026).

19. Collinet, C. & Lecuit, T. Programmed and self-organized flow of information during morphogenesis. Nat. Rev. Mol. Cell Biol. 22, 245–265 (2021).

20. Kinoshita, N., Sasai, N., Misaki, K. & Yonemura, S. Apical accumulation of Rho in the neural plate is important for neural plate cell shape change and neural tube formation. Mol. Biol. Cell 19, 2289–2299 (2008).

21. Kimura-Yoshida, C. et al. Fate specification of neural plate border by canonical Wnt signaling and Grhl3 is crucial for neural tube closure. EBioMedicine 2, 513–527 (2015).

22. Koh, I. & Hagiwara, M. Gradient to sectioning CUBE workflow for the generation and imaging of organoids with localized differentiation. *Commun*. Biol. 6, 299 (2023).

23. Mimura, T. & Inoue, Y. Cell-center-based model for simulating three-dimensional monolayer tissue deformation. J. Theor. Biol. 571, 111560 (2023). doi:10.1016/j.jtbi.2023.111560

24. Mimura, T. & Inoue, Y. Comparison of bending-energy discretization methods for anisotropic meshes in morphogenetic simulations. Transactions of the Japan Society for Computational Engineering and Science 2025, 20250010 (2025). doi:10.11421/jsces.2025.20250010

25. De Marco, P., Merello, E., Mascelli, S. & Capra, V. Current perspectives on the genetic causes of neural tube defects. Neurogenetics 7, 201–221 (2006).

26. Hatta, K., Takagi, S., Fujisawa, H. & Takeichi, M. Spatial and temporal expression pattern of N-cadherin cell adhesion molecules correlated with morphogenetic processes of chicken embryos. Dev. Biol. 120, 215–227 (1987).

27. Dady, A., Blavet, C. & Duband, J.-L. Timing and kinetics of E- to N-cadherin switch during neurulation in the avian embryo. Dev. Dyn. 241, 1333–1349 (2012).

28. MacGowan, J., Cardenas, M. & Williams, M. K. Fold-and-fuse neurulation in zebrafish requires Vangl2. Dev. Biol. 524, 55–68 (2025).

29. Pla, P. & Monsoro-Burq, A. H. The neural border: induction, specification and maturation of the territory that generates neural crest cells. Dev. Biol. 444, S36–S46 (2018).

30. Milet, C. & Monsoro-Burq, A. H. Neural crest induction at the neural plate border in vertebrates. Dev. Biol. 366, 22–33 (2012).

31. Hong, C.-S. & Saint-Jeannet, J.-P. The activity of Pax3 and Zic1 regulates three distinct cell fates at the neural plate border. Mol. Biol. Cell 18, 2192–2202 (2007).

32. Ulloa, F. & Briscoe, J. Morphogens and the control of cell proliferation and patterning in the spinal cord. Cell Cycle 6, 2640–2649 (2007).

33. Le Dréau, G. & Martí, E. Dorsal–ventral patterning of the neural tube: a tale of three signals. Dev. Neurobiol. 72, 1471–1481 (2012).

34. Smith, J. L. & Schoenwolf, G. C. Notochordal induction of cell wedging in the chick neural plate and its role in neural tube formation. J. Exp. Zool. 250, 49–62 (1989).

35. Ybot-Gonzalez, P., Cogram, P., Gerrelli, D. & Copp, A. J. Sonic hedgehog and the molecular regulation of mouse neural tube closure. Development 129, 2507–2517 (2002).

36. Yamada, T., Pfaff, S. L., Edlund, T. & Jessell, T. M. Control of cell pattern in the neural tube: motor neuron induction by diffusible factors from notochord and floor plate. Cell 73, 673– 686 (1993).

37. Yamanaka, Y., et al. Reconstituting human somitogenesis *in vitro*. Nature 614, 509–520 (2023).

38. Gribaudo, S., et al. Self-organizing models of human trunk organogenesis recapitulate spinal cord and spine co-morphogenesis. Nat. Biotechnol., 42, 1243–1253 (2024).

39. Saade M. & Marti E. Early spinal cord development: from neural tube formation to neurogenesis. Nat. Rev. Neurosci 26, 195–213 (2025).

40. Ura T., Honda H., & Takeichi M., Planar Cell Polarity Links Axes of Spatial Dynamics in Neural-Tube Closure. Cell 149, 1084–1097 (2012).

41. Matsuda M., Rozman J., Ostvar S., Kasza K.E. & Sokol SY. Mechanical control of neural plate folding by apical domain alteration. Nat. Commun. 14, 8475 (2023).

42. Mii Y., et al. Planar cell polarity emerges through polarized accumulation of Wnt11. Sci. Adv. 12, eaea0326 (2026).

