## Supplementary Information for "Engineering developmental transitions during human neurulation through modular integration of multimodal spatial cues"

### Methods

We simulated cell invagination and tube formation to examine how morphology depended on slit width and boundary conditions.

##### Cell-centre model and simulation conditions

Three-dimensional epithelial deformation was simulated using a cell-centre model in which tissue shape was described by the positions of cell centres, with the epithelial sheet represented as a triangular network formed by neighbouring cell centres[1,2]. Cell motion, division and rearrangement were based on our previous model[1]. The motion of cell $i$ was governed by the overdamped equation:

$$\gamma \frac{\mathrm{d}\boldsymbol{r}_i}{\mathrm{d}t} = -\nabla_{\boldsymbol{r}_i} E + \boldsymbol{F}_i^{\mathrm{ext}}$$

$$E = E_l + E_A + E_b + E_{\mathrm{rep}} + E_{\mathrm{apical}} + E_{\mathrm{center}} + E_z$$

Here, $E$ is the total energy, $\boldsymbol{r}_i$ is the position of cell $i$, $t$ is time and $\gamma$ is the viscous coefficient. The energy terms denote edge-length elasticity, triangle-area elasticity, bending, repulsion, apical constriction, cell centring and substrate restraint, respectively[1,2]. Bending energy was discretised using the Jülicher formulation[2]. Apical constriction was modelled using two energy terms: a distance-dependent term and a Kantor–Nelson (KN1) angular term incorporating a shifted target dihedral angle[1,2]. The distance term promotes shortening of cell boundary edges, and the angular term favours inward bending. For

$E_{\text{center}}$ and $E_z$, local normals and projected-area weights were held fixed during differentiation. The external force $\boldsymbol{F}_i^{\text{ext}}$ represents Sonic hedgehog (SHH) effects in the tube-formation simulations described below and was zero in the wavelength and invagination analyses.

Cell centres were initially arranged on a triangular lattice. Wavelength and invagination analyses used periodic boundaries in both $x$ and $y$; tube-formation comparisons additionally included free boundaries in both directions. In slit simulations, $x$, $y$ and $z$ denoted the width, longitudinal and height directions, respectively. Height refers to the $z$ coordinate of a cell centre, that is, the out-of-plane displacement of the sheet, and not to cell thickness. Cell division was x-directed: for interior cells, the axis lay in the local tangent plane of the tissue surface and was perpendicular to the global $y$ axis. Boundary cells followed the model's boundary-division rule. Proliferation occurred throughout the tissue, including the substrate-constrained region. Cells initially overlying the slit and their descendants were selected for apical constriction, and substrate restraint towards $z = 0$ was applied only outside the slit. Simulation-specific conditions and shared mechanical and integration parameters are listed in Extended Data Tables 1–3.

Lengths were expressed in units of the rest edge length $l_0$. In the experiments, cell-centre separations in the neural plate region were approximately 10–20 μm immediately after sheet formation, 8 h after seeding. We took one simulation length unit to correspond to 14 µm, a representative value within this observed range.

**Estimation of the characteristic buckling wavelength**

The characteristic wavelength of proliferation-driven buckling was estimated from the dominant peak in the two-dimensional Fourier spectrum of tissue height after 80,000 steps, using a homogeneous-tissue simulation without apical constriction or substrate restraint (Extended Data Table 1). The estimated buckling wavelength, $\lambda = 13.96 l_0$, was used to normalise slit width as $W/\lambda$.

**Mode selection based on wavelength matching**

To estimate the transition between one- and two-wave modes, we assumed that $m = 1$ or $m = 2$ waves fitted within the slit width $W$ and that the selected mode minimised the wavelength mismatch:

$$\delta_m = \left|\frac{W}{m} - \lambda\right|$$

For $\lambda \leq W \leq 2\lambda$, equating the two mismatches gives the transition width $W_c/\lambda = 4/3$.

**Invagination order parameter**

We defined an invagination order parameter, $C$, to quantify the change from a single central trough to two troughs. Twelve equally spaced cross-sections along the periodic $y$ direction were analysed over $-W/2 \leq x \leq W/2$, with the slit centred at $x = 0$.

Profiles were sampled on a uniform $x$ grid with spacing $0.05l_0$. For each cross-section, the straight line joining the slit-edge heights was subtracted. The resulting profile was symmetrised about $x = 0$ and smoothed with a Hann window of width $l_0$ to obtain $h_j(x,t)$. Let $x_{L,j}(t)$ and $x_{R,j}(t)$ be the minimum-height positions on the evaluation grid in $-W/2 \le x < 0$ and $0 < x \le W/2$, respectively, and let $\Delta h_j(t)$ be the maximum-minus-minimum height range of the processed section. We defined:

$$C_j(t) = 1 - \frac{h_j(0,t) - \dfrac{h_j(x_{L,j}(t),t) + h_j(x_{R,j}(t),t)}{2}}{\Delta h_j(t)}$$

The parameter is close to 1 for a single central trough and approaches 0 when the difference between the central height and the mean trough height approaches the full profile height range.

$C$ was evaluated only before an overhang was first detected in any of the 12 cross-sections. Time courses were summarised using the 12 section-wise values and their median. Width comparisons used $C$ evaluated at the last saved time point before the first overhang in each simulation; this time differed between widths. Each width was represented by one simulation. Medians, first-to-third quartile intervals and minimum-to-maximum ranges describe spatial variation among the 12 sections within that simulation.

**Tube formation with contact inhibition**

To represent the effect of SHH, a constant downward force of magnitude $F_0 = 0.20$ was applied to cells in the lowest connected tissue component within $-W/6 < x < W/6$. The targeted component was identified from the current tissue configuration. Tissue fusion was represented by local reconnection of nearby, non-neighbouring surfaces facing one another, subject to geometric and collision checks.

Contact inhibition was determined from the current cell area:

$$A_i = \frac{1}{3}\sum\nolimits_{f \ni i} S_f, \quad A_{\mathrm{cell},0} = \frac{\sqrt{3}}{2} {l_0}^2$$

Here, $S_f$ is the three-dimensional area of an adjacent triangle and $A_{\mathrm{cell},0}$ is the reference cell area. Each triangle's area was shared equally among its three vertices to define the cell areas. When $A_i < rA_{\mathrm{cell},0}$ ($r = 0.90$), division and cell-cycle progression paused; progression resumed from its stored value when the area recovered. The same criterion and reference area were used at free boundaries and after reconnection. Cell-cycle durations were assigned once per generation.

Tube-formation comparisons used periodic sheets with $W = 11l_0$ and $22l_0$, and a sheet with unfixed free boundaries in both directions with $W = 11l_0$. Morphologies were compared at 160,000 steps ($t = 32$), when tube closure in the periodic $W = 11l_0$ control with contact inhibition was confirmed by visual inspection. This common endpoint matched the experimental design, in which all conditions were observed at the control's closure time.

#### References


1 Mimura, T. & Inoue, Y. Cell-center-based model for simulating three-dimensional monolayer tissue deformation. *J. Theor. Biol.* **571**, 111560 (2023). doi:10.1016/j.jtbi.2023.111560

2 Mimura, T. & Inoue, Y. Comparison of bending-energy discretization methods for anisotropic meshes in morphogenetic simulations. *Transactions of the Japan Society for Computational Engineering and Science* **2025,** 20250010 (2025). doi:10.11421/jsces.2025.20250010

**Extended Data Table 1 | Simulation conditions**

| **Condition** | **Invagination simulations** | **Wavelength-estimation simulation** |
|---|---|---|
| Initial lattice ($N_x \times N_y$) | 136 × 64 (8,704 cells) | 350 × 404 (141,400 cells) |
| Periodic lengths ($L_x$, $L_y$) / $l_0$ | (136, 55.4256) | (350, 349.8743) |
| Boundary conditions | Periodic in x and y | Periodic in x and y |
| Initial height / $l_0$ | 0 | U(−0.001, 0.001) |
| Initial cell age | U(0, 20) | U(0, 20) |
| Cell-cycle timescale, mean (s.d.) | 20 (0.2) | 20 (0.2) |
| Division direction | In the local tangent plane, perpendicular to the global y axis | In the local tangent plane, perpendicular to the global y axis |
| Proliferating region | Entire tissue, including substrate-constrained region | Entire tissue |
| Slit width / $l_0$ | 10, 11, 12, 13, 14, 15, 16, 17, 18, 19, 20, 25, 28, 30, 35 | No slit |
| Cells subject to apical constriction | Cells initially at $\|x\| \leq W/2$ and their descendants | None |
| Apical distance coefficient, $k_a$ | 5 | 0 |
| Apical angle coefficient, $k_\theta$ | 10 | 0 |
| Target dihedral angle, $\theta_a$ | π − 0.26 rad | Not applied |
| Substrate coefficient, $k_z$ | 10 | 0 |
| Conditions for substrate restraint | Current $\|x\| > W/2$ and $\|z\| < l_0$ | None |

Lengths and times are in simulation units. U(a, b) denotes a uniform distribution on (a, b).

**Extended Data Table 2 | Shared mechanical and integration parameters**

| **Parameter** | **Symbol** | **Value** |
|---|---|---|
| Rest edge length | $l_0$ | 1 |
| Reference triangle area | $A_{\triangle,0}$ | 0.433 |
| Edge-length stiffness | $k_l$ | 2 |
| Area stiffness | $k_A$ | 10 |
| Bending coefficient | $\kappa$ | 8.660254 |
| Spontaneous curvature | $H_0$ | 0 |
| Repulsive coefficient | $k_{\mathrm{rep}}$ | 10 |
| Repulsive cutoff distance | $r_c$ | 0.9 |
| Centring-force coefficient | $k_c$ | 10 |
| Viscous coefficient | $\gamma$ | 0.25 |
| Time step | $\Delta t$ | 0.0002 |
| Time-integration method | — | Explicit Euler |

Values are reported in simulation units. Energy formulations were based on ref. 1, with Jülicher bending and the KN1 angular term based on ref. 2. Apical constriction and substrate coefficients differed between simulations as listed in Extended Data Tables 1 and 3.

**Extended Data Table 3 | Tube-formation comparisons**

| **Condition** | **Slit width, $W/l_0$** | **Boundary conditions** |
|---|---|---|
| Control | 11 | Periodic in x and y |
| Wide slit | 22 | Periodic in x and y |
| Free boundary | 11 | Free in x and y |

All conditions used the initial lattice size, cell-cycle timescale, and apical and substrate parameters of the invagination simulations in Extended Data Table 1, with the shared mechanical parameters in Extended Data Table 2. Contact-inhibition ratio $r = 0.90$; downward force per targeted cell $F_0 = 0.20$. All conditions used x-directed division and were compared at 160,000 steps.
